# The histologic phenotype of lung cancers may be driven by transcriptomic features rather than genomic characteristics

**DOI:** 10.1101/2021.01.01.425056

**Authors:** Ming Tang, Hussein A Abbas, Marcelo Vailati Negrao, Maheshwari Ramineni, Xin Hu, Junya Fujimoto, Alexdrandre Reuben, Susan Varghese, Jianhua Zhang, Jun Li, Chi-Wan Chow, Xizeng Mao, Xingzhi Song, Won-chul Lee, Jia Wu, Latasha Little, Curtis Gumbs, Carmen Behrens, Cesar Moran, Annikka Weissferdt, J.Jack Lee, Boris Sepesi, Stephen Swisher, John V. Heymach, Ignacio I. Wistuba, P. Andrew Futreal, Neda Kalhor, Jianjun Zhang

## Abstract

Histology plays an essential role in therapeutic decision-making for lung cancer patients. However, the molecular determinants of lung cancer histology are largely unknown. We conducted whole-exome sequencing(WES) and microarray profiling on 19 micro-dissected tumor regions of different histologic subtypes from 9 patients with lung cancers of mixed histology. A median of 68.9% of point mutations and 83% of copy number aberrations were shared between different histologic components within the same tumors. Furthermore, different histologic components within the tumors demonstrated similar subclonal architecture. On the other hand, transcriptomic profiling revealed shared pathways between the same histologic subtypes from different patients, which was supported by the analyses of the transcriptomic data from 141 cell lines and 343 lung cancers of different histologic subtypes. These data suggest that histology of lung cancers may be determined at the transcriptomic level rather than the genomic level.

## Introduction

Lung cancer is the leading cause of cancer death in the United States with an estimated 228,820 new cases and 135,520 deaths expected in 2020^1^. Histopathology continues to play an essential role in prognosis and choosing appropriate treatment^2^. Largely determined by morphology, primary lung cancers are histologically classified into small cell lung cancers (SCLC) and non-small cell lung cancers (NSCLC) and the latter include adenocarcinoma (LUAD), squamous cell (LUSC), and large-cell neuroendocrine (LCNEC) as the main histologic subtypes. However, consensus histologic confirmation can sometimes be challenging and therefore impacts optimal treatment choices^3,4^.

The molecular mechanisms determining the tumor histology are unknown. Previous studies revealed that tumors from different patients or even multiple independent primary lung cancers within the same patients can have identical morphology yet shared no mutations^5^, while there can be a morphologic difference in different regions within the same tumors that share the majority of mutations^6^. These findings suggest that morphology may not be primarily determined by genomic features.

About 5% of primary lung cancers can present with a mixed histologic pattern, where additional components with distinct histologic types present within the same tumors, often referred to as combined or mixed histology^7,8^. Tumors with mixed histology provide a unique opportunity to study the molecular basis for histology determination as different histologic components share the same clinical and genetic backgrounds, and exposure history. There have been a few studies on lung cancers of mixed histology, most of which focused on the genomic changes of adenosquamous lung cancers. The majority of these studies revealed shared driver mutations between different histologic components^8–14^. These findings are overall in line with the prior hypothesis that genomic changes were not the main determinants of histology. However, these studies only covered hotspot driver mutations or small gene panels while mutations of other genes with essential biological functions and other genomic alterations such as somatic copy number alterations (SCNA) were not investigated. Thus the relationship between genomic alterations and histology was not fully addressed.

In the current study, we leveraged three unique datasets to fill this void: 1) whole-exome sequencing (WES) and transcriptomic data from 19 micro-dissected tumor regions of different histology from 9 primary lung cancer patients with mixed histologic patterns including 6 LUAD, 6 LCNEC, 3 SCLC, 3 LUSC, and one poorly differentiated NSCLC-NOS; 2) transcriptomic data from 141 cell lines of different histologic subtypes from the cancer cell line encyclopedia (CCLE)^15^ including 14 LCNEC, 57 LUAD, 48 SCLC, and 22 LUSC; 3) transcriptomic data from a total of 343 patients including 14 LCNEC, 273 LUAD, 9 SCLC and 47 LUSC with lung cancers of different histologic subtypes^16,17^.

## Results

### Patients characteristics

The clinicopathologic characteristics of the 9 patients with lung cancers of mixed histology are summarized in Table 1. The median age at diagnosis with lung cancer was 67 years (range 47-79 years). All patients were current (3/9) or former (6/9) smokers. Eight patients had two different distinct histologic subtypes, while one patient had 3 different histologic subtypes (Table 1). Different histologic components of each tumor of mixed histology were manually micro-dissected, which resulted in 19 different tumor tissues including 6 LUAD, 6 LCNEC, 3 SCLC, 3 LUSC, and one poorly differentiated NSCLC-NOS that were subjected to WES and microarray RNA profiling. The most common combination of mixed histology was LCNEC-LUAD in 4/9 patients, followed by LCNEC-LUSC and SCLC-LUAD subtypes in 2/9 patients each, and one patient had SCLC-LUSC subtypes.

### Shared mutations across different patients and distinct histologic subtypes

We first investigated whether the mutations overlapped between different histologic components within the same tumors and whether there were particular mutations shared across the same histologies from different patients. Overall, different histologic components from the same tumors shared the majority of mutations (**Fig. 1** and **Supplementary Fig. 1 a-i**). The percentage of shared mutations within the same tumors ranged from 12.1% to 98.4% with a median of 68.9%, similar to that between different regions within the same tumors of the same histology (68.9% vs 72%, p=0.46, Wilcoxon rank-sum test)^6^. Of note, except Pa35, who had 12.1% shared mutations between the SCLC and LUAD histologic subtypes, all other patients had >40% mutations shared amongst distinct histologies. On the other hand, the same histologic subtypes across different patients barely shared any mutations. Specifically, only 1 intronic mutation in *RABL6* (chr9:139731607:A:G) was shared between LUAD components of Pa31 and Pa35 as well as the SCLC components of the same two patients, whereas no mutations were shared among the 6 LCNEC or 3 LUSC histologic subtypes consistent with previous findings ^6^ suggesting somatic mutations may not be the primary determinant of histology in these tumors.

**Fig 1:**
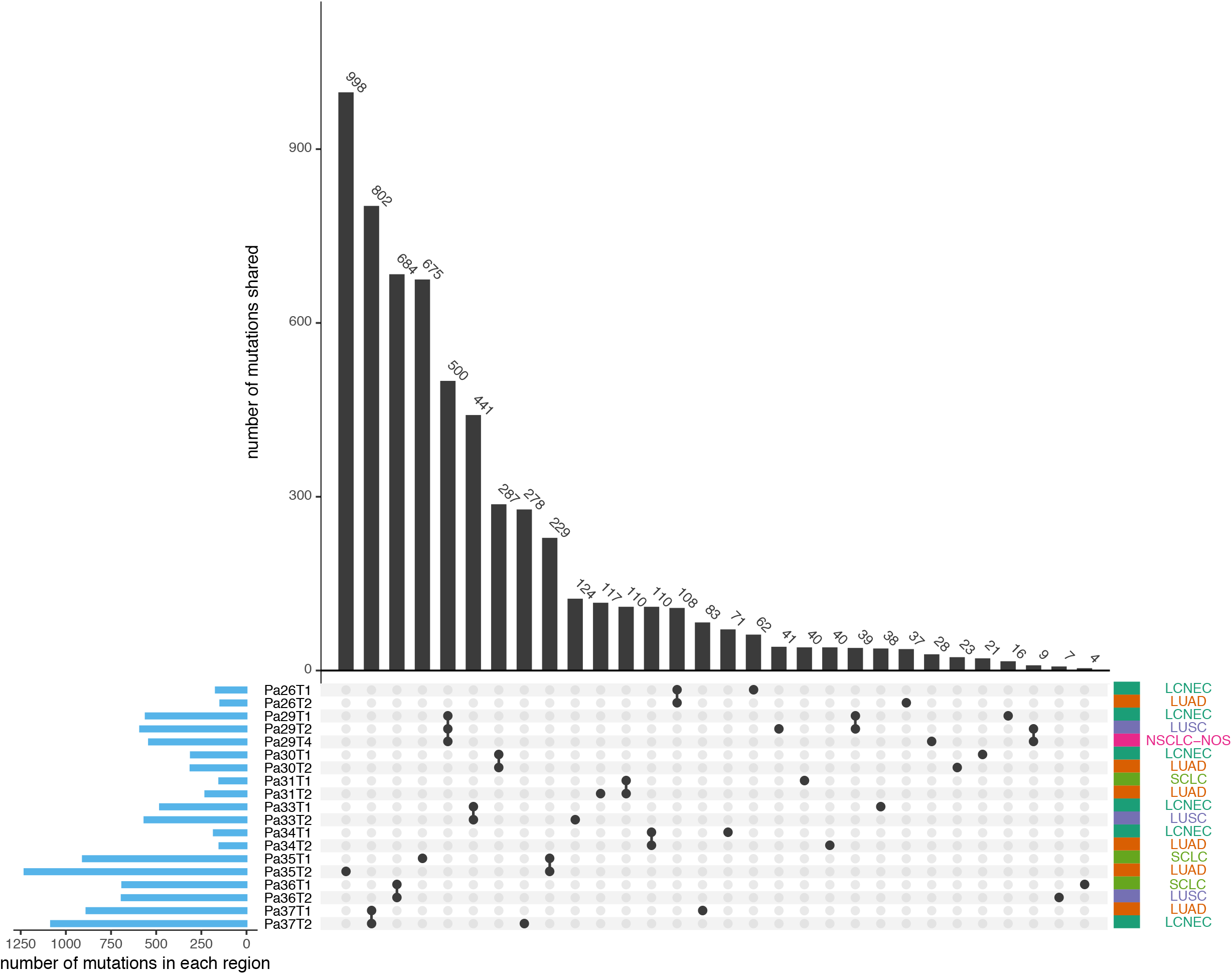
Overlapping somatic mutations across different histologic subtypes within the same patient. The upset plot demonstrates the shared mutations across samples. Blue bars in the y-axis represent the total number of mutations in each sample. Black bars in the x-axis represent the number of mutations shared across samples connected by the black dots in the body of the plot.

### Similar mutational processes between different histologic subtypes within the same tumors

It is well known that different cancer types have distinct mutational signatures^18^ suggesting different mutational processes in play reflecting different genetic backgrounds and exposure etiologies associated with different cancer types. To understand whether the mutational processes are histology-specific in these lung tumors of mixed histology in the context of identical genetic background and exposure history, we calculated the mutational spectrum and mutational signatures in each histologic component. Overall, a similar mutational spectrum was observed between different histologic components within the same tumors (**Fig. 2a**). We next calculated the contribution of 30 signatures of mutational processes in cancer^18^ (**Fig. 2b-c**). Not surprisingly, Signature 4 (associated with smoking and tobacco carcinogenesis) was the most dominant in 7 of 9 patients consistent with their smoking history (**Fig. 2c**). Two exceptions were patients Pa35 and Pa26, who were both former light smokers with a 2.5 and 5 pack-year smoking history, respectively and both quit >20 years ago. Other common signatures in this cohort of tumors included Signature 1 (associated with spontaneous deamination of 5-methylcytosine), Signatures 2 and 13 (associated with APOBEC-mediated mutagenesis), and Signature 16 (etiology-unknown). Similar to the mutation spectrum, the mutational signatures were also overall similar between different histologic components within the same tumors, while none of the mutational signatures enriched in certain histologic components was shared across different patients. Taken together, these data suggest that mutational processes were not histology-specific, but rather patient-specific, likely determined by the particular exposure history and host factors in each patient.

**Fig 2:**
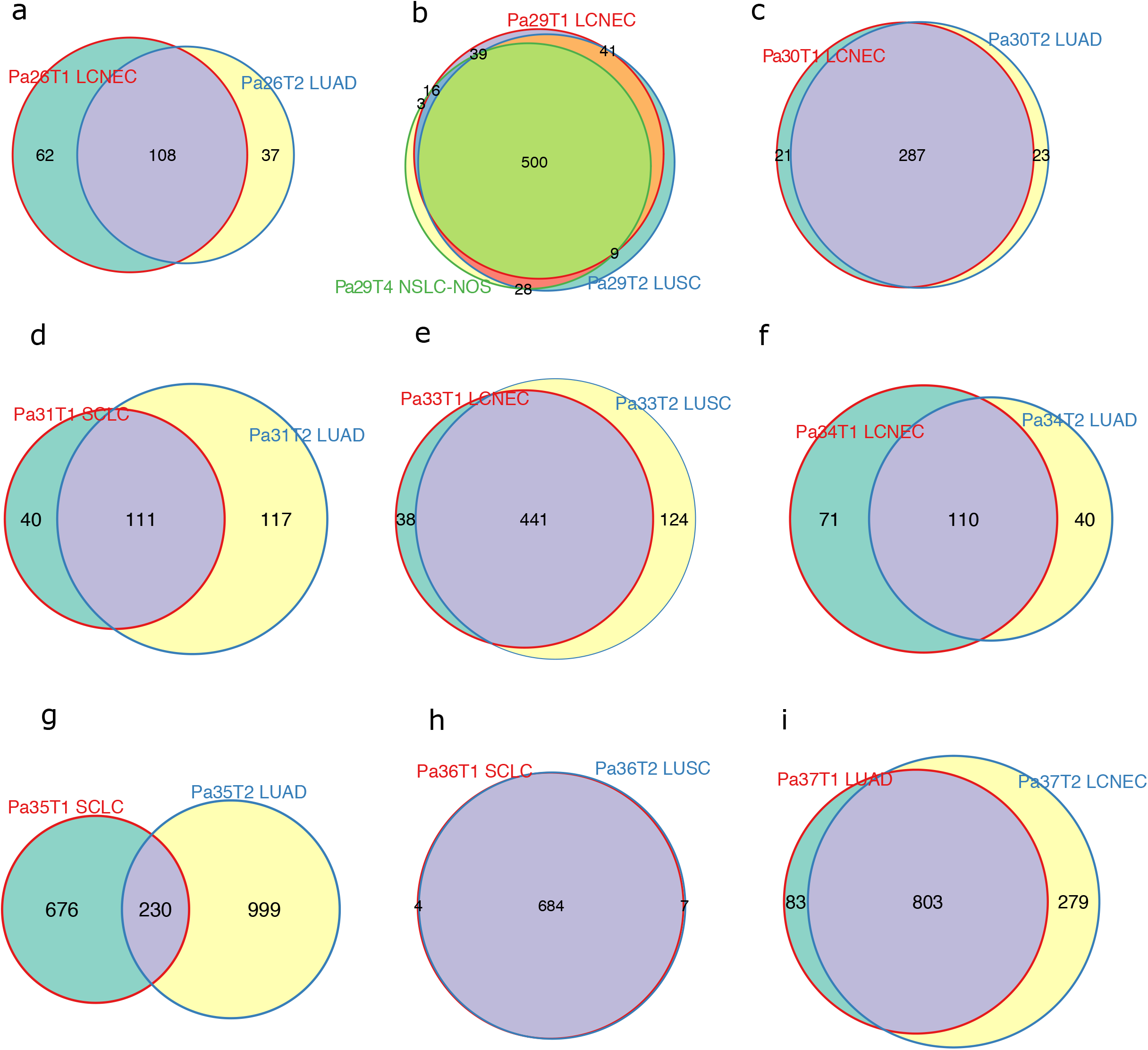
Mutational spectrums and signatures are similar across different histologic subtypes within the same patient. (a) Bar plots represent the mutational spectrum decomposed by trinucleotide context. (b) Heatmap of the contribution of the 30-COSMIC mutation signatures in each sample. (c) Stacked barplot for the contribution of the top 10 mutation signatures in each sample.

### Similar subclonal architecture between different histologic components

We next inferred cancer cell fractions (CCF) of all somatic mutations using PyClone^19^ to determine the subclonal architecture in each histologic component. Overall, the subclonal architecture was similar between different histologic components within the same tumors. A substantial proportion of clonal mutations^20,21^, were shared across different histologic components of the same tumors and only a small proportion of clonal mutations were private (**Fig. 3a-k**). Specifically, among the shared mutations, an average of 54.6% (ranging 16% − 96.5%) were clonal, while only 10.7% (ranging 0.14% − 35.8%) private mutations were clonal. Taken together, these data support that those different histologic components within the same tumors were derived from the same progenitor cell and that the divergence of distinct histologic components was a relatively late molecular event.

**Fig 3:**
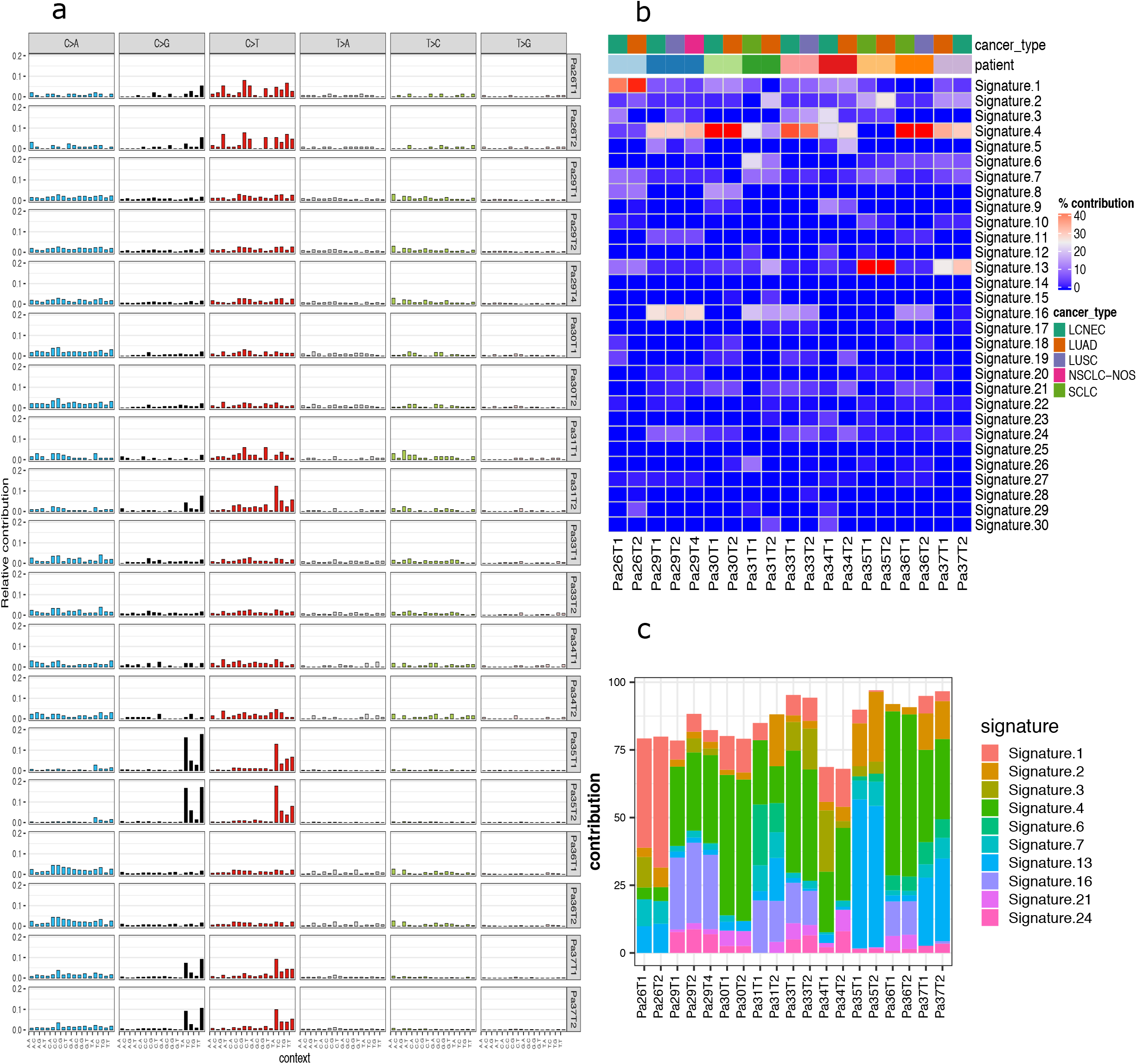
Clonality analysis revealed shared clonal mutations between different histologies within the same patients. (a-k) Scatter plots of the cellular prevalence of somatic mutations calculated by PyClone for the two histological components within the same patient. Mutations were clustered by PyClone and mutations of the same cluster were labeled with the same color. Labeled genes represent canonical cancer gene mutations.

### Similar somatic copy number aberration profiles between different histologic subtypes within the same tumors

SCNA is another key feature of human malignancies that could potentially impact the expression of large groups of genes. We next delineated the genome-wide SCNA profiles. As shown in **Fig. 4a** and **4b**, the overall SCNA profiles were similar between different histologic components within the same patients, while drastically different among different patients. Furthermore, we quantified SCNA events using a gene-based SCNA analysis algorithm^22^ for exome sequencing data that allows comparing the SCNAs between different samples to identify shared and unique SCNA events between different histologic components within the same tumors. To minimize the impact of tumor purity on SCNA analysis, we obtained purity-adjusted log2 copy number ratios for each tumor in this study (see Methods for details). On average, 83% of SCNA events (ranging from 54.7% to 99.1%) were shared between different histologic components within the same tumors suggesting the majority of SCNA events were early molecular events before the separation of different histologic components. No particular SCNAs were found to be enriched in certain histologic subtypes. Furthermore, compared to the intratumor heterogeneity (ITH) dataset from the TRACERx study ^25^, at the gene level, the extent of shared SCNA landscape between different histologic components was comparable to that between spatially separated tumor regions within the same NSCLC tumors of the same histology (83% in mixing histology cohort vs 72% in TRACERx cohort, *p*=0.25, Wilcoxon rank-sum test).

**Fig 4:**
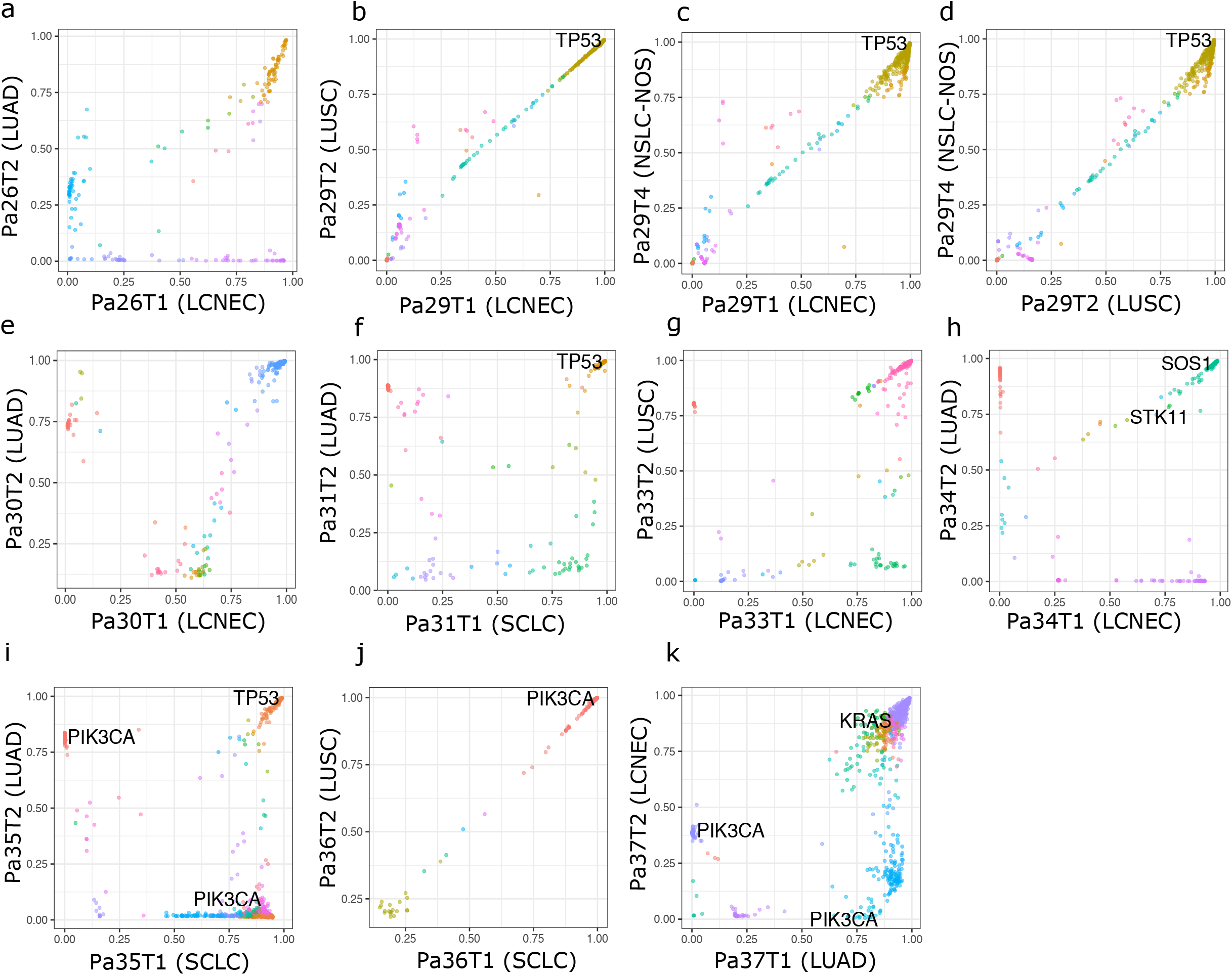
Somatic copy number aberration (SCNA) analysis demonstrated similar copy-number changes between different histologic subtypes within the same patient. (a) IGV screenshot of genome-wide SCNA profile for each sample. (b) Heatmap of the correlation of SCNA at gene-level. (c) Heatmap of copy number changes from canonical cancer genes of the COSMIC database.

### The majority of cancer gene alterations occurred before the divergence of different histologic components of the same tumors

Cancer gene mutations are known to determine distinct molecular subsets of lung cancers with unique clinical presentation and cancer biology and certain cancer gene mutations are even considered pathognomonic for certain histologic subtypes^23^. Therefore, we investigated whether specific cancer gene mutations could determine different histologic patterns in these tumors of mixed histology. A total of 11 canonical cancer gene mutations, defined as nonsynonymous mutations identical to those previously reported in known cancer genes^24,25^ or truncating mutations in known tumor suppressor genes, were identified in these 19 specimens and 10 of the 11 canonical cancer gene mutations were clonal in each histologic component except for *PIK3CA* (p.E545K) in Pa37 LCNEC component that was subclonal (**Fig. 3**). Furthermore, 7 of the 11 cancer gene mutations were shared between different histologic components within the same tumor, while 4 were private. Interestingly, in patient Pa35, a *PIK3CA* p.M1043I mutation was shared between the SCLC (CCF = 0.9) and LUAD (CCF = 0.09) components while a *PIK3CA* p.E542K was only detected in the LUAD component (CCF = 0.8) (**Fig. 3i**). Similarly, in Pa37, a *PIK3CA* p.E545K was identified in both LUAD (CCF = 0.009) and LCNEC (CCF = 0.4) components, while a *PIK3CA* p.H1047R was private to the LUAD component (CCF = 0.85) (**Fig. 3k**). These findings are reminiscent of heterogeneity studies in kidney ^26^ and lung cancers^5,27,28^, where different mutations in the same cancer genes were identified in different regions within the same tumors or different independent primary tumors within the same patients. These results imply convergent evolution that even with an identical genetic background and environmental exposure, the evolution of different cancer cell subclones can be driven by distinct molecular events, with possible genetic constraints around certain genes or pathways (*PIK3CA* in case of patient Pa35 and Pa37) that are pivotal for cancer evolution.

Furthermore, we estimated copy number gains of oncogenes and copy losses of tumor suppressor genes (TSG) in this cohort of tumors of mixed histology based on the COSMIC database^24^(**Fig. 4c**). A total of 11 copy number gains of 5 oncogenes and 129 copy number losses of 27 TSGs were detected in this cohort of tumors of mixed histology. Similar to cancer gene point mutations, 53.8% of SCNA in oncogenes and TSGs were shared within the same patients. These data suggested that the cancer gene mutations and copy number changes were early molecular events acquired before the divergence of different histologic subtypes and maybe not the major mechanisms determining the histologic fate of cancer cells in lung cancers of mixed histology.

### Specific transcriptomic patterns may be associated with specific histologic subtypes

As the histology of lung cancers did not appear to be determined by genomic aberrations, we next sought to explore whether the cell fate is determined at the transcriptomic level. We first performed the gene expression profiling of the same tumor regions of distinct histologic subtypes to investigate whether transcriptomic signatures could differentiate histological subtypes. By principal component analysis (PCA), the normal lung tissues were separated from the tumor samples highlighting the distinct transcriptomic changes associated with malignant cells (**Fig. 5a**). Tumor specimens of different histologic subtypes from the same patients overall clustered together although there was a small cluster of LUAD samples from different patients clustered close to each other (**Fig. 5a**). In unsupervised hierarchical clustering, different histologic components within the same tumors also tended to cluster together highlighting substantial inter-patient heterogeneity. On the other hand, 8 of the 19 specimens were clustered with specimens from a different patient, significantly more common than that of different tumor regions within the same tumors of same histology, where 2 of 35 specimens were clustered with a different patient (p=0.001 by Chi-Square test)^29^. Among these 8 specimens, 4 LUAD specimens (Pa26T2, Pa30T2, Pa31T2, and Pa37T1) were clustered together, while Pa35T1 (LCNEC) clustered with Pa37T2 (SCLC) and P30T1 (LCNEC) clustered with P31T1 (SCLC) (**Fig. 5b**) − both LCNEC and SCLC are considered as neuroendocrine tumors sharing many biological and clinical features^30^. Similarly, the LCNEC components of patients Pa26 and Pa29 were clustered together. Taken together, these data suggested that in the background of patient-specific gene expression profiles, there may be histology-specific transcriptomic features, associated with distinct histological phenotypes.

**Fig 5:**
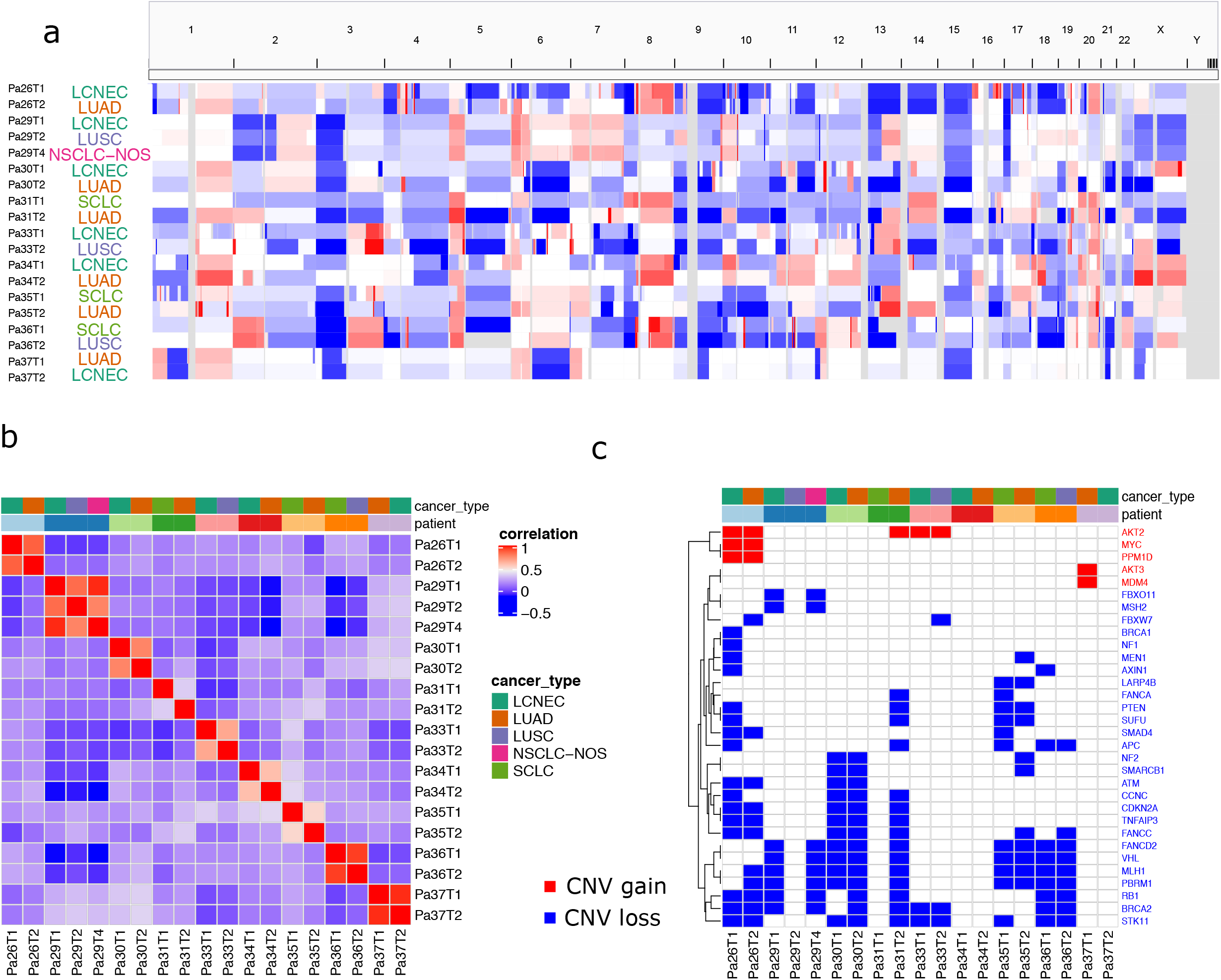
Gene expression profile revealed some extent of similarity of the same histologic components across different patients. (a) Principal component analysis (PCA) of all histologic subtypes based on gene expression data. (b) Heatmap of the top 500 most variable genes across the samples clustered by both genes and the samples. (c) Commonly up-regulated and down-regulated pathways comparing SCLC with LUAD across public datasets and in-house dataset. (d) Commonly up-regulated and down-regulated pathways comparing LCNEC with LUAD across public datasets and in-house dataset.

### Histology-specific pathways shared with independent cohorts

To further understand the transcriptomic features associated with different histologies, we evaluated if any Hallmark pathways^31^ were enriched in different histologic subtypes. To identify histology-specific pathways, we looked specifically at overlapping pathways in the histologic comparison pairs in different patients that had the same direction of enrichment (either positive or negative). The most concordant pattern was noted in Pa31 and Pa35 with SCLC versus LUAD, whereas 3 pathways were upregulated and 9 pathways were down-regulated in SCLC components compared to LUAD components (**Fig. 5c**). Interestingly, the 3 up-regulated pathways in SCLC (E2F_Target, G2M_checkpoint, and MYC_target) were associated with cell proliferation while 6 of the 9 down-regulated pathways in SCLC components (IL2, complement, INFG, INFA, TNFA, and inflammatory response) were associated with inflammatory/immune response. In the LCNEC versus LUAD comparisons, there were no pathways with consistent enrichment in all 4 patients. However, compared to LUAD, MYC, G2M, and E2F pathways were up-regulated in LCNEC components in 3/4, ¾, and 2/4 and patients, respectively, while interferon-alpha and interferon-gamma responses were down-regulated in LCNEC components in 2/4 and 2/4 patients, respectively (**Supplementary Table 2**).

To validate these findings, we analyzed the transcriptomic data from another 3 different cohorts: two previously published large cohorts of primary lung cancers by Karlsson et al^17^, which encompassed 126 primary lung cancers (83 LUAD, 26 LUSC, 3 SCLC, and 14 LCNEC) and by Bhattacharjee et al^16^ with 217 lung cancer patients (190 LUAD, 21 LUSC, and 6 SCLC), as well as 141 cell lines (57 LUAD, 22 LUSC, 48 SCLC, and 14 LCNEC) from CCLE database^32^. Using the same approach for data from tumors of mixed histology, we identified enriched pathways by comparing LCNEC versus LUAD, LCNEC versus LUSC, SCLC versus LUAD, and SCLC versus LUSC of each cohort respectively (**Supplementary Table 2**). We next focused on the pathways that were 1) identified in at least 2 patients from our mixed histology cohort and 2) validated by at least two of the 3 datasets (Karlsson cohort, Bhattacharjee cohort, and CCLE). With these criteria, SCLC versus LUAD comparison demonstrated the most consistent pattern with cell proliferation-related pathways upregulated and inflammatory/immune response pathways downregulated in SCLC (p. adj<0.05) (**Fig. 5c**). Also, for LCNEC versus LUAD histology pathway analysis, there was significant positive enrichment for cell cycle G/M cell cycle checkpoint and MYC targets for Pa30, Pa34, Pa37, and in the Karlson dataset (p.adj <0.05) (**Fig. 5d**).

## Discussion

In the immuno-oncology era, histological subtype remains to play essential roles in determining the optimal treatment for lung cancer patients^34–36^ For example, surgical resection is the main treatment modality for localized NSCLC, while SCLC is usually treated with chemotherapy and radiation even at localized stage^33^. In the metastatic setting, the chemotherapy regimens are also different for different histologic subtypes.

Currently, the mechanisms underlying histologic cell fate are unknown. Understanding the molecular determinants of histology may provide novel insights to understand the different responses to various treatment regimens and to more effectively leverage histology to guide lung cancer management. Although large-scale studies such as in TCGA have demonstrated that genomic features are largely distinct between different lung cancer histologic subtypes^23,37,38^, genomic alterations do not always agree with histologic subtypes. Targetable genomic alterations such as *EGFR* mutations and *ALK/ROS1* translocations that are pathognomonic for LUAD have been reported in some LUSC patients and SCLC patients^39,40^, suggesting that the histology is not primarily determined by genomic features. However, these analyses are complicated by the distinct genetic background and exposure history in different cancer patients.

Cancers of mixed histology provide a unique opportunity to identify the molecular features associated with different histologic components in the setting of identical genetic background and exposure history. Among the mixed histology lung tumors, the adenosquamous is the most frequently studied subtype while other mixed histology subtypes were rarely investigated. In the current study, we focused on lung cancers of non-adenosquamous subtypes and applied WES and gene expression microarray with the intent to depict the comprehensive molecular bases of histology. Analysis of WES data from 9 patients with mixed histology demonstrated that different histological components within the same tumors shared a large proportion of identical point mutations, which is consistent with previous studies in adenosquamous subtypes by cancer gene panel sequencing^7–11^. In addition to more comprehensive point mutation data, WES also allowed us to compare different histologic components regarding the SCNA profiles, which demonstrated that different histologic components from the same tumors share the majority of SCNA events. Furthermore, different histologic components from the same tumors also demonstrated overall similar subclonal architecture and canonical cancer gene alterations. Taken together, these data suggest that different histologic components were derived from the same progenitor cells and that the divergence of distinct histologic components was a relatively late molecular event conferring inter-histologic heterogeneity. Thus, the histologic subtype was not primarily determined by genomic alterations.

There is ample evidence that gene expression profiling can inform lung cancer histology^16,17,41^. Our transcriptomic profiling from histologic subtypes in tumors of the same patient allowed decoupling of the effect of the patient’s genetic background and exposures in influencing the transcriptomic signatures. Unlike the similar genomic landscape between different histologic components, intra-tumor heterogeneity of transcriptomic profiles between different histologic components was significantly higher than spatially separated regions from tumors of the same histology. A substantial proportion of tumor regions clustered more closely together with tumor regions of the same histology from different patients, significantly more common than that in different tumor regions of the same histology^29^. Pathway analysis demonstrated common pathways between different histologic components across different patients, which were further supported by integrative analysis from cell lines and larger cohorts of patient datasets. These were mostly accentuated between SCLC and LUAD as well as LCNEC and LUAD. Compared to LUAD components, SCLC and LCNEC tumors, both of which are high-grade neuroendocrine carcinomas, demonstrated up-regulation of pathways associated with cell proliferation including G2M, E2F, and MYC consistent with the high proliferative nature of SCLC and LCNEC^42^. Of particular interest, 6 of 9 down-regulated pathways in SCLC were inflammatory/immune pathways in line with reported cold immune microenvironment and inferior response to immunotherapy in SCLC^43^. These results also suggest histology-specific modulation of the tumor microenvironment even within the same tumors with the same genetic background and exposure.

In summary, we sought to provide novel insights to dissect the molecular basis for the histologic determination by multi-omics analysis of 3 unique datasets: lung cancers of mixed histology providing a unique opportunity to identify the molecular features associated with different histologic components in the setting of identical genetic background and exposure history; CCLE cell lines of different histology allowing analyzing pure epithelial cancer cells without confounding effect from stromal components; and large cohorts of human lung cancers of different histologic subtypes. Our analysis demonstrated that the different histologic components from the same patients share the majority of point mutations, SCNA, and cancer gene alterations suggesting a shared cell of origin and indicated that histology may not be determined at the genomic level. On the other hand, although essentially no genomic mutations were shared, different tumor regions of the same histology across different patients tended to be more closely clustered based on transcriptomic profiles. Pathway analysis revealed important pathways encircled for certain histologies, which were validated by CCLE cell lines and two large cohorts of human lung cancers. These data suggested that histology of lung cancers may be determined at the transcriptomic level although the exact mechanisms of gene expression regulation remain to be determined. These intriguing findings have to be validated on larger cohorts of tumors of mixed histology and by functional analyses in future studies.

## Methods

### Sample Collection and Processing

Patients with mixed histology lung cancer were included in this study after confirmation with two independent pathologists. Unstained slides were microdissected after delineating the different regions of histologic components and then extracted for RNA and DNA. A written informed consent that was approved by the internal review board of the University of Texas M D Anderson Cancer Center was obtained. The study was conducted in accordance with the Declaration of Helsinki.

### Whole-exome sequencing

DNA was extracted using the QIAamp DNA FFPE Tissue Kit (QIAGEN) and the resulting genomic DNA was sheared into 300–400□bp segments and subjected to library preparation for whole-exome sequencing using KAPA library prep (Kapa Biosystems) with the Agilent SureSelect Human All Exon V4 kit according to the manufacturer’s instructions. Paired-end multiplex sequencing of DNA samples was performed on the Illumina HiSeq 2000 sequencing platform.

### Somatic mutation calling and overlapping mutations

The whole-exome sequencing raw FASTQ files were aligned using bwa-mem. Mutations were called using mutect^44^ and Lancet following GATK best practice (www.broadinstitute.org/gatk/guide/best-practices.php) for duplicate removal, indel realignment, and base recalibration. Lancet^45^ was used for SNV and indel calling using localized colored de Bruijn graph. For SNVs, only those which were called by more than one caller or called in more than one sample from the same patient were retained. For all mutations, we recovered the raw allelic counts from the bam file if it occurred in one of the different histologic subtypes from the same patient. The process was implemented as a Snakemake pipeline and can be found at https://gitlab.com/tangming2005/snakemake_DNAseq_pipeline/tree/multiRG. The number of overlapping mutations across all samples were plotted in an UpSet plot^46^ and Venn diagrams.

### Clonal architecture analysis

A high-quality list of SNVs was combined from all samples from the same patient and the allelic counts for those positions were obtained using bam-readcount (https://github.com/genome/bam-readcount) Copy number variations and tumor purity were obtained from sequenza^47^, and the mutation allelic counts were analyzed with PyClone for clonality analysis^19^. PyClone was run with 10,000 iterations and a burn-in of 1,000 as suggested by the authors.

### Mutational signature and spectrum analysis

Mutation signatures and spectrum analysis were analyzed by Bioconductor package MutationalPatterns^48^ with 30 COSMIC signatures following the standard workflow.

### Somatic Copy number analysis (SCNA)

Copy number analysis was carried out using Sequenza^47^. Both copy number and tumor purity were inferred by Sequenza. Since the signal to noise ratio of SCNA could be reduced in the samples with lower tumor purity, we obtained purity-adjusted log2 ratios by log2((original copy ratio-1)/purity+1)^49^. The segment files were visualized in IGV^50^. We then used the log2 thresholds of log2(4/2) and log2(1/2) to determine whether a gene is gained or lost focusing only on cancer genes that have shown to have copy number changes in the COSMIC database. The matrices of log_2_ ratio or binarized copy number status for all genes and cancer genes, respectively, across all samples, were clustered using hierarchical clustering and plotted in a heatmap using ComplexHeatmap^51^.

### In-house microarray and public microarray/RNAseq data analysis

The in-house clariom.s.human microarray data were analyzed using Bioconductor packages Oligo^52^, pd.clariom.s.human, and limma^53^ following standard workflow. GSE94601 microarray data were downloaded using GEOquery^54^ and analyzed by the limma package. The Bhattacharjee et al microarray data were downloaded from http://portals.broadinstitute.org/cgi-bin/cancer/publications/view/62 and analyzed using the affy^55^ and limma package. The CCLE lung cancer RNAseq count data were downloaded from the Broad CCLE data portal and processed using DESeq2^56^. Gene set enrichment analysis using Hallmark dataset was carried out using fgsea Bioconductor package^57^ and the genes are pre-ranked by (signed log2FoldChange) * −log_10_(p-value) for all the public datasets. For the in-house microarray data, we computed the fold change between distinct histologies within the same patient and rank the genes by the fold change.

## Data availability and code availability

The whole-exome sequencing data have been deposited at the European Bioinformatics Institute European Genome–phenome Archive (EGA) (accession number: pending) through controlled access. To protect patient privacy, interested researchers need to apply via data access committee (DAC), which will grant all reasonable requests. And source data are provided with this study. All other data may be found within the main manuscript or supplementary information or available from the authors upon request. Public and in-house microarray ExpressionSet objects can be found at https://osf.io/gxc4r/. The code used to generate the figures can be found at https://github.com/crazyhottommy/mixed_histology_lung_cancer.

## Acknowledgments

This study was supported by the National Cancer Institute of the National Institute of Health Research Project Grant (R01CA234629-01), the AACR-Johnson & Johnson Lung Cancer Innovation Science Grant (18-90-52-ZHAN), the MD Anderson Physician Scientist Program, the MD Anderson Lung Cancer Moon Shot Program, TJ Martell Foundation Award, Sabin Family Foundation Award, Duncan Family Institute Cancer Prevention Research Seed Funding Program, the Cancer Prevention and Research Institute of Texas Multi-Investigator Research Award grant (RP160668) and the UT Lung Specialized Programs of Research Excellence Grant (P50CA70907), Cancer Prevention and Research Institute of Texas (CPRIT) grant RP150079. HAA was supported in part by the T32 NIH fellowship.

## Competing Interests

Dr. Zhang reports research funding from Merck, Johnson and Johnson, and consultant fees from BMS, Johnson and Johnson, AstraZeneca, Geneplus, OrigMed, Innovent outside the submitted work. Dr. Heymach reports research funding from AstraZeneca, GlaxoSmithKline, and Spectrum; consultant fees from AstraZeneca, Boehringer Ingelheim, Bristol-Myers Squibb, Catalyst, EMD Serono, Foundation Medicine, Hengrui Therapeutics, Genentech, GSK, Guardant Health, Eli Lilly, Merck, Novartis, Pfizer, Roche, Sanofi, Seattle Genetics, Spectrum, and Takeda; licensing fees from Spectrum. Dr. Kadara reports funding from Johnson and Johnson and Janssen Pharmaceuticals. Dr. Sepesi reports consultant fees from BMS. The other authors declare neither financial nor non-financial interests in the submitted work.

---

**Supplementary Fig 1.**
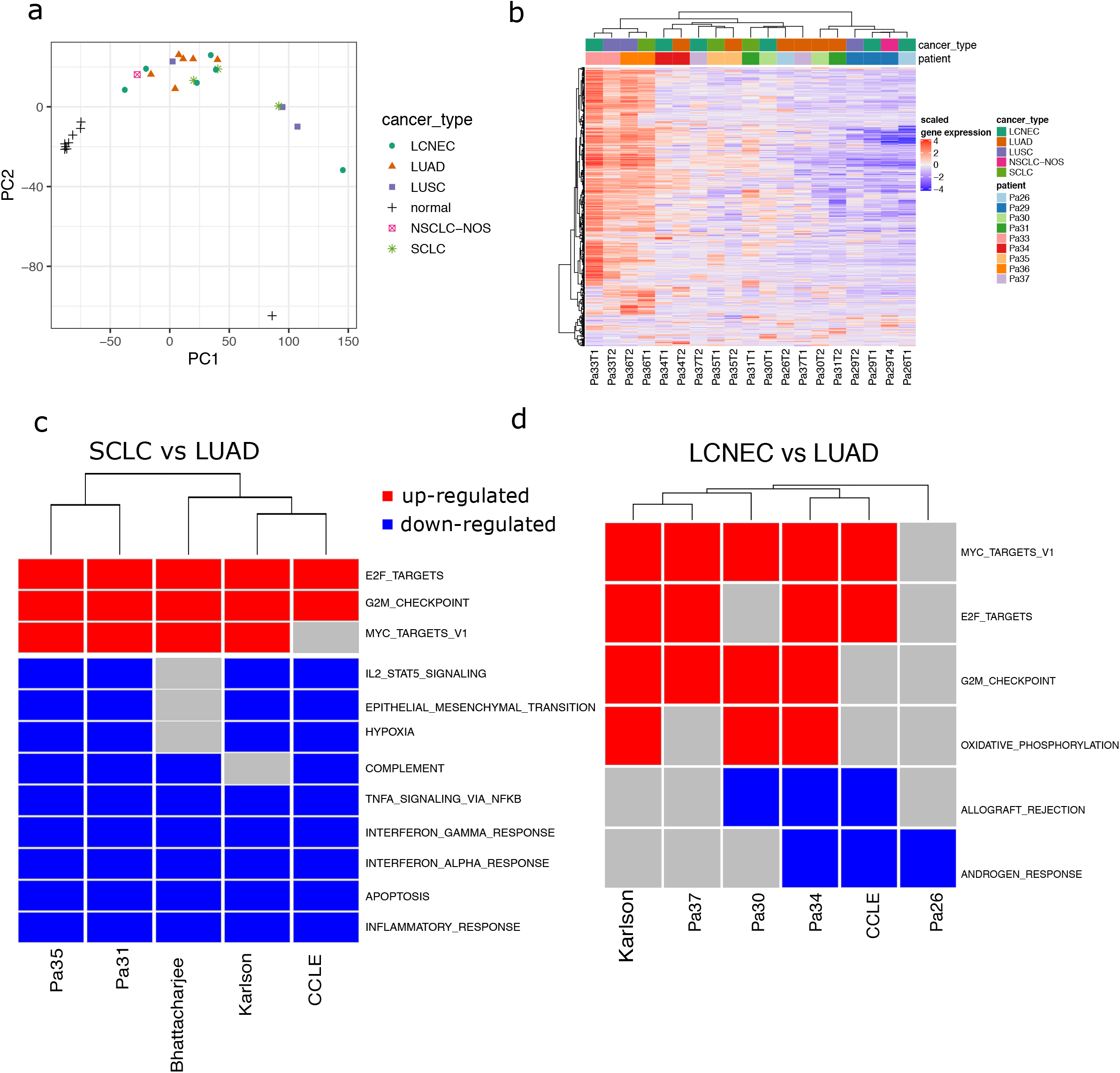
(a-i) Venn-diagram showing the number of overlapping mutations in each patient across different histological components.

